# A Robust Method to Estimate the Largest Lyapunov Exponent of Noisy Signals: A Revision to the Rosenstein’s Algorithm

**DOI:** 10.1101/381111

**Authors:** Sina Mehdizadeh

## Abstract

**Aim:** This study proposed a revision to the Rosenstein’s method of numerical calculation of largest Lyapunov exponent (LyE) to make it more robust to noise.

**Methods:** To this aim, the effect of increasing number of initial neighboring points on the LyE value was investigated and compared to the values obtained by filtering the time series. Both simulated (Lorenz and passive dynamic walker) and experimental (human walking) time series were used to calculate LyE. The number of initial neighbors used to calculate LyE for all time series was 1 (the original Rosenstein’s method), 2, 3, 4, 5, 10, 15, 20, 25, and 30 data points.

**Results:** The results demonstrated that the LyE graph reached a plateau at the 15-point neighboring condition inferring that the LyE values calculated using at least 15 neighboring points were consistent and reliable.

**Conclusion:** The proposed method could be used to calculate LyE more reliably in experimental time series acquired from biological systems where noise is omnipresent.

## 1. Introduction

The last two decades have seen a growing trend towards adopting the nonlinear measure of largest Lyapunov exponent (LyE) to quantify structure variability in experimental time series examples of which are human movement [1–4], heart rate [5], electromyography [6]. The LyE measures the exponential rate of divergence of neighboring trajectories of the state space constructed by data acquired from experiment [7]. Rosenstein’s method [8] is the most common numerical method of calculating LyE from experimental data. A recent review on gait demonstrated that 79% of the studies on walking used Rosenstein’s method to calculate LyE [9].

One of the main challenges encountered with the calculation of LyE using experimental time series is the measurement noise [9–11]. The presence of such noise could have adverse effects on the calculated LyE since it increases the possibility of picking false neighbors in the state space [11, 12] and thus computation of unreliable LyE values [13]. Studies using Rosenstein’s method reported different values of error in the presence of noise. For instance, in the original study of Rosenstein et al., [8], up to 80% error was found for LyE value of different attractors in the presence of noise. Using Lorenz and Rossler attractors, Rispens, Pijnappels, van Dieёn, van Schooten, Beek and Daffertshofer [14] also demonstrated that the percent error of the LyE values was in the range of 10–100%. In a recent study, Mehdizadeh and Sanjari [13] also reported approximately 60% error for LyE values associated with the simplest passive dynamic walker under different noise levels. Although Rosenstein, Collins and De Luca [8] suggested filtering the signal to rectify noise problem, filtering may lead to loss of information contained in the signal [15]. Therefore, much uncertainty still exists about the proper method of reducing the adverse effect of noise on the LyE.

In order to understand the effect of noise, it is fundamental to know how Rosenstein’s method computes LyE. To calculate LyE, first, an appropriate state space is constructed using the experimental time series to completely embed the system’s dynamics. Then, the distance between two neighboring points (i.e., initial conditions) of the state space is measured and tracked over the entire evolution time of the system. This process is repeated for and averaged over all points of the state space. The neighboring point here is the nearest point (amongst all points of the state space) to the reference point (Figure 1). However, when the time series is contaminated by noise, it is possible that the nearest point be a noise point rather than a real data point. More specifically, the probability of the selected nearest point being a noise point is 50% when only a single point is selected as the nearest neighbor. As a consequence of selecting the noise as the nearest point, the divergence of the neighboring trajectories might not be exponential. This is evident in the divergence graph of noisy time series where the graph does not resemble a stereotypical divergence graph (see Figure 2 in Mehdizadeh and Sanjari [13]). Therefore, the slope of the linear part of divergence graph (i.e. the rate of divergence of trajectories) will not indicate LyE.

**Figure 1:**
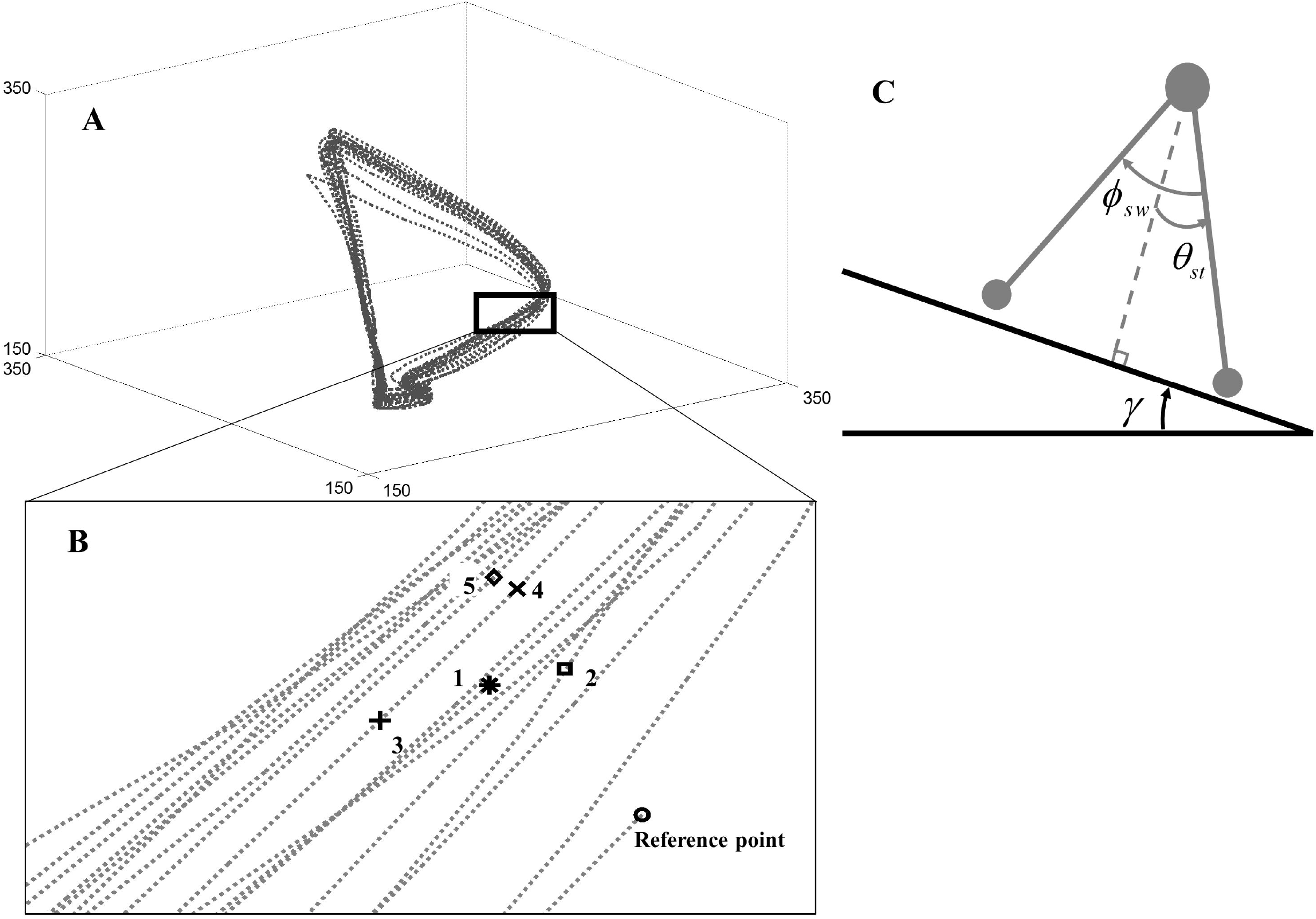
A) 3-dimensional representation of a state space constructed from experimental data. Note that the actual dimension of a state space might be greater than three. However, only up to three dimensions could be shown visually. B) the expanded view of the rectangular area in the state space to show the separation of an exemplar reference point (○) and the first (*), second (□), third (+), fourth (×), and fifth (◊) neighboring points. The distance to the reference point was 39.31, 44.82, 73.63, 79.65, and 83.10 mm for the first to fifth neighboring points, respectively. Note that the distance between each neighboring point and the reference points is an n-dimensional Euclidean distance. Therefore, the distances in the 2-dimensional figure might not necessarily be the actual distance. The number of each neighboring point is also depicted in the figure. Only the first point is used to calculate the divergence at each time step in the original Rosenstein’s method. C) Schematic representation of the passive dynamic walker model. See the text for the details of the model.

One possible solution to overcome the noise problem could be increasing the number of neighboring points (initial conditions). Hence, by selecting 2 neighboring points for instance, the probability of both points being noise reduces to 25%; by selecting 3 points, the probability reduces to 12.5% and so on. It should be noted that this modification does not violate the assumptions of the original Rosenstein’s algorithm because it calculates the divergence at each time increment as the average distance among multiple nearest points at that time increment (Figure 1; see the methods section for details). Kantz[12] used a similar approach to modify Wolf’s[16] algorithm in computing LyE. However, no attempts have been made for the Rosenstein’s method to make it more robust to noise and this is the first time that this issue is being tackled.

The first aim of this study was therefore to investigate the effect of increasing the number of initial neighboring points on the LyE value in the presence of noise. Furthermore, since Rosenstein, Collins and De Luca [8] suggested filtering to rectify noise, a second aim of the study was to compare the LyE calculated using the proposed method to that of using filtering method. Time series associated with both mathematical systems (Lorenz and simplest passive dynamic walker) and biological systems (human walking recorded using motion capture and accelerometer) were used to calculate LyE.

## 2. Methods

### 2.1. The new algorithm

The approach proposed here is a revision to Rosenstein et al. (1993), algorithm in that the nearest neighbors of the reference point are more than one and the initial separation (*d_j_ave_*(0)) is the average of Euclidean distance of all points to the reference point (equation 1):

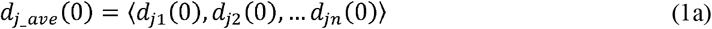

where 〈.〉 denotes averaging and *dj_n_*(0) is the distance between *n*^th^ nearest neighbor, *X_jn_* to the reference point 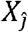 (equation 1b):

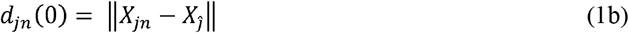

where ||.|| denotes Euclidean distance. In practice, *d_jn_*(0) can be found by calculating the distance of all state space data points to the reference point and sorting them from lowest to highest. The first point in the sorted series is the nearest point (the only point used in the original Rosenstein’s method), the second point is the second nearest point and so on. The important point here to consider is that all nearest points should be located at a distance greater than the mean period of the time series (i.e. should not be located at the same trajectory on which the reference point is located).

From this point on, the algorithm follows the original algorithm proposed by Rosenstein, Collins and De Luca [8]. That is, the largest finite-time Lyapunov Exponent (*λ_max_*) could be determined using equation (1c):

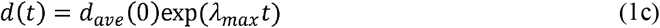

where *d*(*t*) is the average distance between neighboring points at time *t*, and the initial separation of the neighboring points is represented by *d_ave_*(0). According to equation (1c), for the *j^th^* pairs of neighboring points in state space we have:

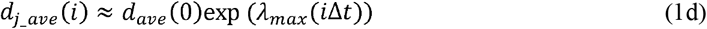

Taking the logarithm from both sides of (1d) results in:

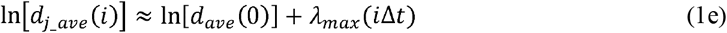

*λ_max_* is determined using a linear fit to the following curve::

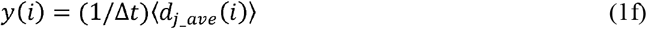

where 〈*d_j_ave_*(*i*)〉 denotes the average over all pairs of.

### 2.2. Mathematical models

#### 2.2.1. Lorenz model

The differential equation of the Lorenz system is represented in equation (2). The parameters of the system were according to Rosenstein, Collins and De Luca [8], and were *σ* = 16.0, *R* = 45.92, and *b* = 4.0. These equations were integrated numerically to obtain the time series associated with *x, y* and *z*

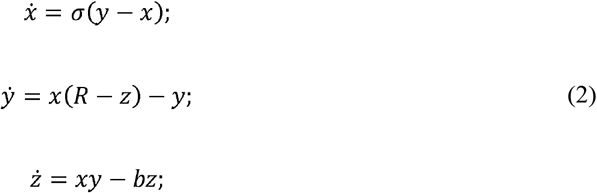

#### 2.2.2. Passive dynamic walker model

The simplest passive dynamic walker (PDW) model has been widely adopted in the studies of bipedal locomotion[13, 17]. The model comprised of two massless legs connected by a frictionless hinge joint at the hip. It has point masses at the hip and the feet (Figure 1). The model assumes that the feet masses are significantly small compared to the hip mass. When walking down a slope of angle *γ*, the stance leg rotates as an inverted pendulum until the swing leg contacts the ground. At the contact point, the swing leg becomes the new stance leg and the stance leg becomes the new swing leg for the next step. It has been shown that the model exhibits a period-1 limit-cycle behavior for *γ*≤0.019 rad [17]. The motion of the model is described by two second-order differential equations:

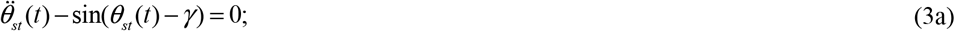

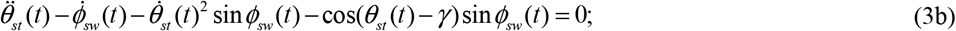

where *θ_st_* is the angle of the stance leg with respect to the perpendicular to *γ* and *φ_sw_* is the angle of the swing leg with respect to the stance limb, *t* is time and all derivatives are with respect to time. For this model, the swing foot contact (when the legs switch) occurs when the swing leg angle is twice the stance leg angle:

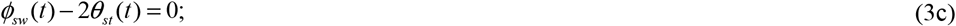

At this point, switching of the two legs occurs using equation (3d)[17]:

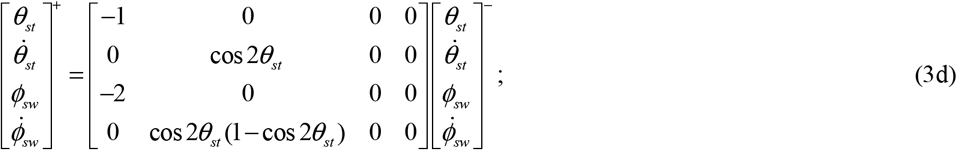

where the “+” superscript indicates the system state just after the foot contact and the “-” superscript indicates the system state just before the foot contact. Walking data were generated by integrating equations (3a) and (3b) considering switching specified in Equations (3c) and (3d). For our study, γwas set at 0.009 rad.

### 2.3. Experiments

Time series from human walking were used as the examples of biological time series. Since in the studies on human movement, LyE is quantified for time series obtained from either motion capture system or accelerometers, this study also acquired data of human walking using both systems. A young male participant (age = 32 yrs, mass = 63.0 kg, and height = 1.75 m) walked on a motorized treadmill (Bertec, Columbus, USA) on his preferred walking speed (the participant signed an informed consent before participating in the study). Fifteen Qualisys Oqus (Qualisys AB, Gothenburg, Sweden) calibrated cameras recorded three-dimensional coordinate of reflective markers attached to the skin of bony landmarks of sacrum, and left ankle (lateral malleolus). Reconstruction and labeling were performed using Qualisys QTM software (Qualisys AB, Gothenburg, Sweden). In addition, two calibrated Noraxon Myomotion inertial sensors (Noraxon, Scottsdale, USA) capable of recording 3-dimensional accelerations (range: 16g) were attached to the sacrum landmark and also on the lateral side of the left shank’s distal end just above lateral malleolus. Noraxon MR 3.10 (Noraxon, Scottsdale, USA) software was used to acquire acceleration data. Three trials of 3-minute waking was done while position and acceleration data being simultaneously recorded at the sampling frequency of 100 Hz.

### 2.4. Data analysis

For all simulated and experimental time series, 50 consecutive cycles from the middle of the time series were chosen and rescaled to 5000 data points for further analysis (approximately 100 points per cycle). This was done because studies have reported at least 50 consecutive cycles are needed for a reliable estimation of LyE[13]. For simulated time series, five levels of Gaussian white noise (GWN) (signal to noise ratio (SNR) = 80-0dB with 20 dB steps) were added to the time series of *x* and *θ* for Lorenz and PDW systems, respectively. The choice of the amplitude of the noise was so that to be in the range of previous studies[8, 13, 14]. GWN is used as it simulates noise commonly encountered in experimental conditions[11]. In addition, to compare the effect of filtering with the proposed revised method, experimental time series were filtered using a 4^th^ order lowpass Butterworth filter with a cutoff frequency determined using residual method [13, 18]. A state space with time delay of 10 and embedding dimension of 5 was constructed using the average mutual information[19] and global false nearest neighbors[20] methods, respectively. The number of initial neighbors used to calculate LyE for all time series was 1 (the original Rosenstein’s method), 2, 3, 4, 5, 10, 15, 20, 25, and 30. Furthermore, LyE was calculated for the time series of *x* (Lorenz system), *θ* (PDW system), sacrum and ankle 3-dimensional acceleration (accelerometer data) and 3-dimensional velocity (motion capture data; marker position data were differentiated to remove non-stationarities). Finally, the %Difference per point was calculated as the % difference between the LyE values of two consecutive neighboring condition.

## 3. Results

### 3.1 Simulated time series

The LyE values calculated for Lorenz system in different noise levels and with different number of neighboring points are gathered in Table 1. The associated %Difference is also shown in Figure 2. The %Difference reaches a plateau at the 15-point neighboring condition (less than 1% difference) onwards for all noise levels. Table 2 and Figure 2 also indicate the LyE values, and %Difference for PDW, respectively. Here the %Difference also reaches a plateau on the 15-point condition (also less than 1% difference) and remains in this region for all noise levels.

**Figure 2:**
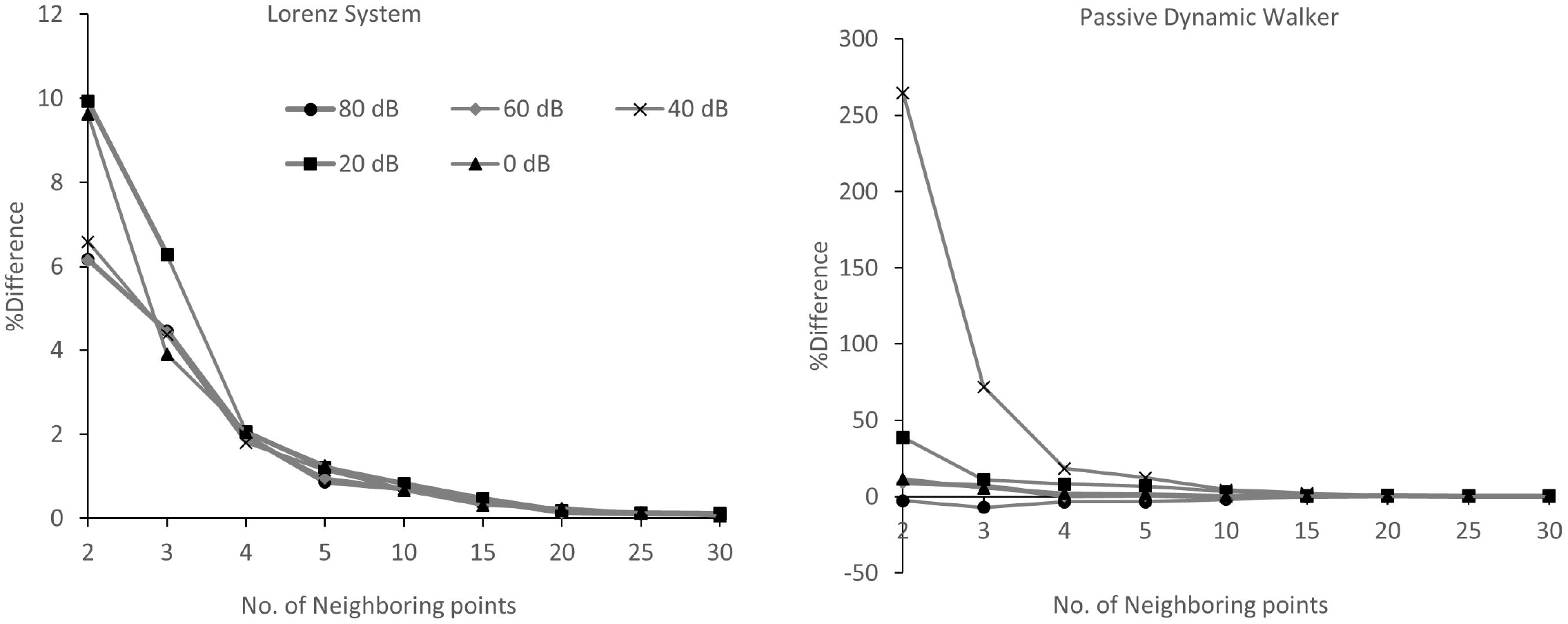
The %Difference calculated for the Lorenz (left) and passive dynamic walker (right) systems under different noise conditions and with different number of neighboring points.

**Table 1:**
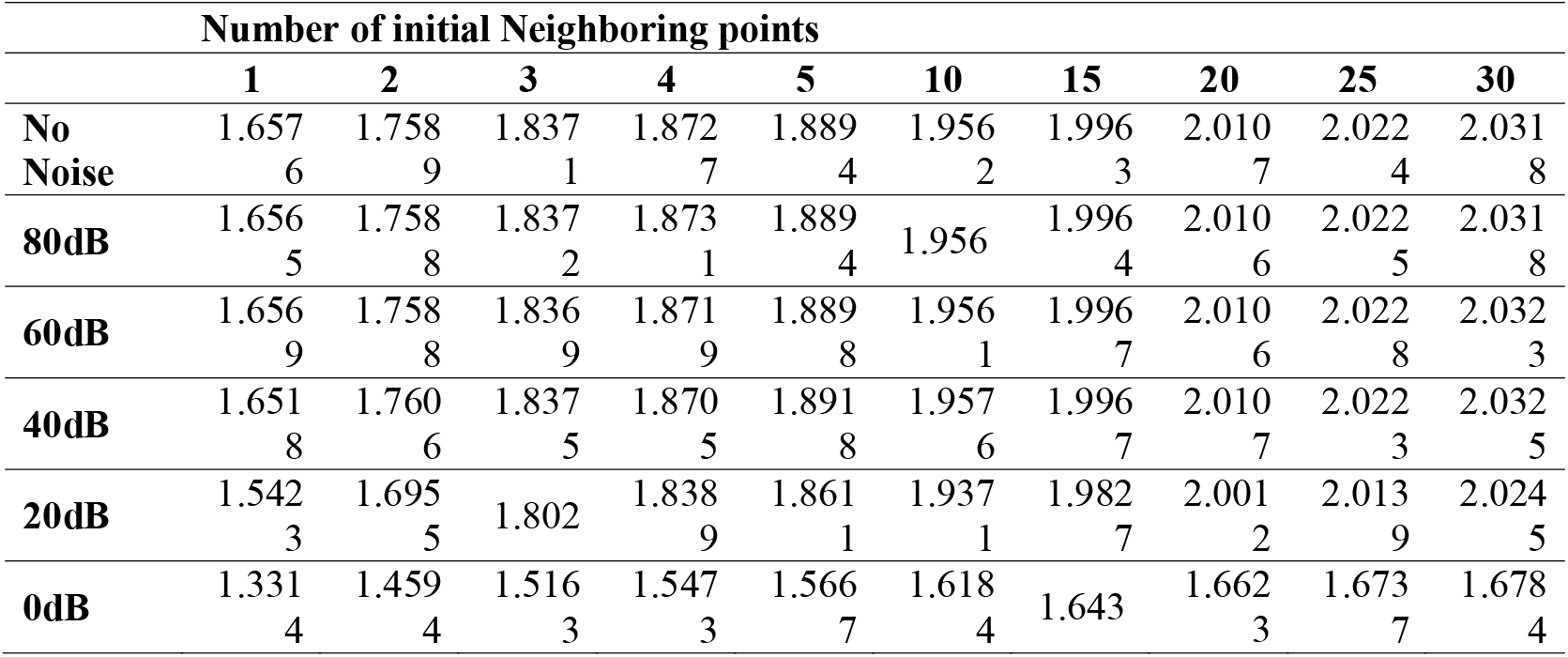
The largest Lyapunov exponent (LyE) calculated for noise-free and noisy Lorenz system using different number of neighboring points. The 1-point neighboring condition is equivalent to the original Rosenstein’s method.

|  | Number of initial Neighboring points |  |  |  |  |  |  |  |  |  |
| --- | --- | --- | --- | --- | --- | --- | --- | --- | --- | --- |
|  | 1 | 2 | 3 | 4 | 5 | 10 | 15 | 20 | 25 | 30 |
| **No Noise** | 1.657<br>6 | 1.758<br>9 | 1.837<br>1 | 1.872<br>7 | 1.889<br>4 | 1.956<br>2 | 1.996<br>3 | 2.010<br>7 | 2.022<br>4 | 2.031<br>8 |
| **80dB** | 1.656<br>5 | 1.758<br>8 | 1.837<br>2 | 1.873<br>1 | 1.889<br>4 | 1.956 | 1.996<br>4 | 2.010<br>6 | 2.022<br>5 | 2.031<br>8 |
| **60dB** | 1.656<br>9 | 1.758<br>8 | 1.836<br>9 | 1.871<br>9 | 1.889<br>8 | 1.956<br>1 | 1.996<br>7 | 2.010<br>6 | 2.022<br>8 | 2.032<br>3 |
| **40dB** | 1.651<br>8 | 1.760<br>6 | 1.837<br>5 | 1.870<br>5 | 1.891<br>8 | 1.957<br>6 | 1.996<br>7 | 2.010<br>7 | 2.022<br>3 | 2.032<br>5 |
| **20dB** | 1.542<br>3 | 1.695<br>5 | 1.802 | 1.838<br>9 | 1.861<br>1 | 1.937<br>1 | 1.982<br>7 | 2.001<br>2 | 2.013<br>9 | 2.024<br>5 |
| **0dB** | 1.331<br>4 | 1.459<br>4 | 1.516<br>3 | 1.547<br>3 | 1.566<br>7 | 1.618<br>4 | 1.643 | 1.662<br>3 | 1.673<br>7 | 1.678<br>4 |

**Table 2:** The largest Lyapunov exponent (LyE) calculated for noise-free and noisy passive dynamic walker using different number of neighboring points. The 1-point neighboring condition is equivalent to the original Rosenstein’s method

|  | **Number of initial Neighboring points** |  |  |  |  |  |  |  |  |  |
| --- | --- | --- | --- | --- | --- | --- | --- | --- | --- | --- |
|  | **1** | **2** | **3** | **4** | **5** | **10** | **15** | **20** | **25** | **30** |
| **No Noise** | 0.1932 | 0.1580 | 0.0980 | 0.0690 | 0.0573 | 0.0212 | 0.0189 | 0.0308 | 0.0501 | 0.0780 |
| **80dB** | 0.0994 | 0.0970 | 0.0890 | 0.0868 | 0.0838 | 0.0767 | 0.0763 | 0.0771 | 0.0783 | 0.0830 |
| **60dB** | 0.0681 | 0.0740 | 0.0796 | 0.0802 | 0.0809 | 0.0802 | 0.0803 | 0.0826 | 0.0839 | 0.0850 |
| **40dB** | 0.0034 | 0.0140 | 0.0230 | 0.0252 | 0.0283 | 0.0345 | 0.0382 | 0.0389 | 0.0398 | 0.0440 |
| **20dB** | -0.0294 | -0.0180 | -0.0160 | -0.0147 | -0.0137 | -0.0114 | -0.0111 | -0.0106 | -0.0103 | -0.0100 |
| **0dB** | -0.0530 | -0.0469 | -0.0443 | -0.0433 | -0.0425 | -0.0419 | -0.0412 | -0.0409 | -0.0406 | -0.0404 |

### 3.2. Experimental time series

The LyE values for sacrum and ankle using motion capture system and the associated %Difference are depicted in Table 3 and Figure 3, respectively. For both sacrum and ankle, the graph of %Difference reaches a plateau and the %Difference fell below 1% at the 15-point condition for all directions. Moreover, the %Difference is negative in all graphs except for ankle vertical direction. The results for Acceleration data is similar to motion capture (Table 4 and Figure 4). That is, the graphs reach a plateau and the %Difference fell under 1% at the 15-point neighboring condition. Finally, filtering the signals with the residual frequency (Table 5) resulted in higher LyE values compared to non-filtered signals.

**Figure 3:**
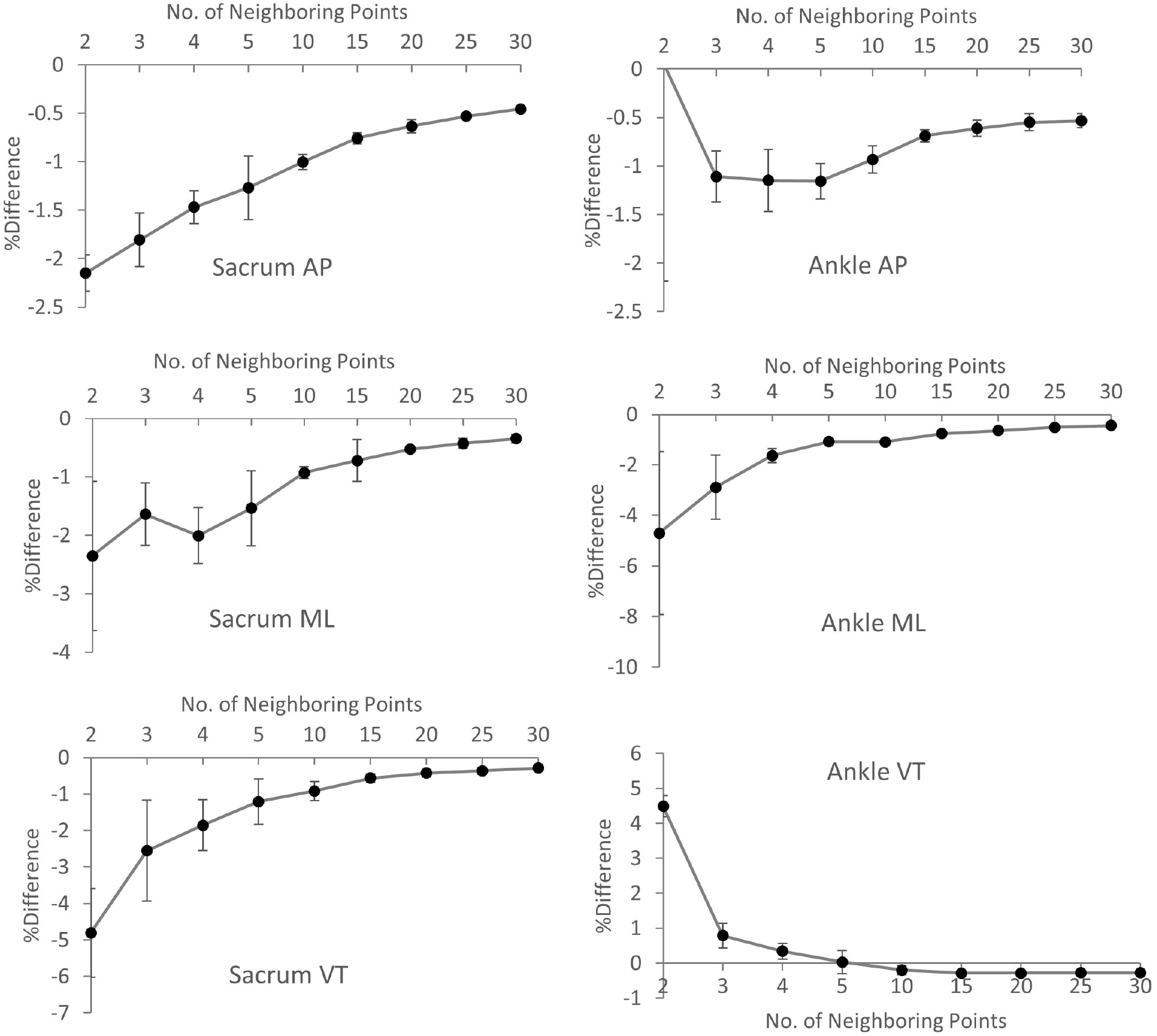
The %Difference calculated for sacrum (left) and ankle (right) velocity time series in anterior-posterior (AP, top), medio-lateral (ML, middle), and vertical (VT, bottom) directions. The motion capture system was used to record markers’ position. The 1-point neighboring condition is equivalent to the original Rosenstein’s method.

**Table 3:** Average (standard deviation) of the largest Lyapunov exponent (LyE) calculated for the velocity time series of sacrum and ankle acquired from the motion capture system. The values are averaged over three trials. The 1-point neighboring condition is equivalent to the original Rosenstein’s method.

| Number of initial Neighboring points |  |  |  |  |  |  |  |  |  |  |
| --- | --- | --- | --- | --- | --- | --- | --- | --- | --- | --- |
|  | 1 | 2 | 3 | 4 | 5 | 10 | 15 | 20 | 25 | 30 |
| **Sacrum** |  |  |  |  |  |  |  |  |  |  |
| **AP** | 0.4959<br>(0.0446<br>) | 0.4853<br>(0.0443<br>) | 0.4764<br>(0.0425<br>) | 0.4695<br>(0.0425<br>) | 0.4636<br>(0.0430<br>) | 0.4405<br>(0.0422<br>) | 0.4239<br>(0.0415<br>) | 0.4106<br>(0.0412<br>) | 0.3997<br>(0.0404<br>) | 0.3906<br>(0.0398<br>) |
| **ML** | 0.2372<br>(0.0527<br>) | 0.2321<br>(0.0536<br>) | 0.2285<br>(0.0536<br>) | 0.2241<br>(0.0533<br>) | 0.2209<br>(0.0535<br>) | 0.2105<br>(0.0503<br>) | 0.2035(<br>0.0511) | 0.1982<br>(0.0501<br>) | 0.1941<br>(0.0496<br>) | 0.1909<br>(0.0493<br>) |
| **VT** | 0.2856<br>(0.0612<br>) | 0.2722<br>(0.0601<br>) | 0.2655<br>(0.0611<br>) | 0.2609<br>(0.0619<br>) | 0.2579<br>6<br>(0.0625<br>) | 0.2466<br>(0.0630<br>) | 0.2399<br>(0.0629<br>) | 0.2350<br>(0.0625<br>) | 0.2309<br>(0.0626<br>) | 0.2277<br>(0.0624<br>) |
| **Ankle** |  |  |  |  |  |  |  |  |  |  |
| **AP** | 0.8779<br>(0.0389<br>) | 0.8787<br>(0.0550<br>) | 0.8690<br>(0.0544<br>) | 0.8591<br>(0.0558<br>) | 0.8492<br>(0.0566<br>) | 0.8099<br>(0.0593<br>) | 0.7819<br>(0.0560<br>) | 0.7580<br>(0.0553<br>) | 0.7371<br>(0.0543<br>) | 0.7175<br>(0.0551<br>) |
| **ML** | 0.3546<br>(0.0450<br>) | 0.3390<br>(0.0510<br>) | 0.3296<br>(0.0525<br>) | 0.3243<br>(0.0523<br>) | 0.3208<br>(0.0517<br>) | 0.3033<br>(0.0499<br>) | 0.2919<br>(0.0492<br>) | 0.2826<br>(0.0483<br>) | 0.2755<br>(0.0481<br>) | 0.2694<br>(0.0474<br>) |
| **VT** | 0.7620<br>(0.0093<br>) | 0.7962<br>(0.0113<br>) | 0.8025<br>(0.0142<br>) | 0.8053<br>(0.0153<br>) | 0.8056<br>(0.0175<br>) | 0.7975<br>(0.0145<br>) | 0.7860<br>(0.0148<br>) | 0.7749<br>(0.0143<br>) | 0.7640<br>(0.0154<br>) | 0.7534<br>(0.0168<br>) |

**Table 4:**
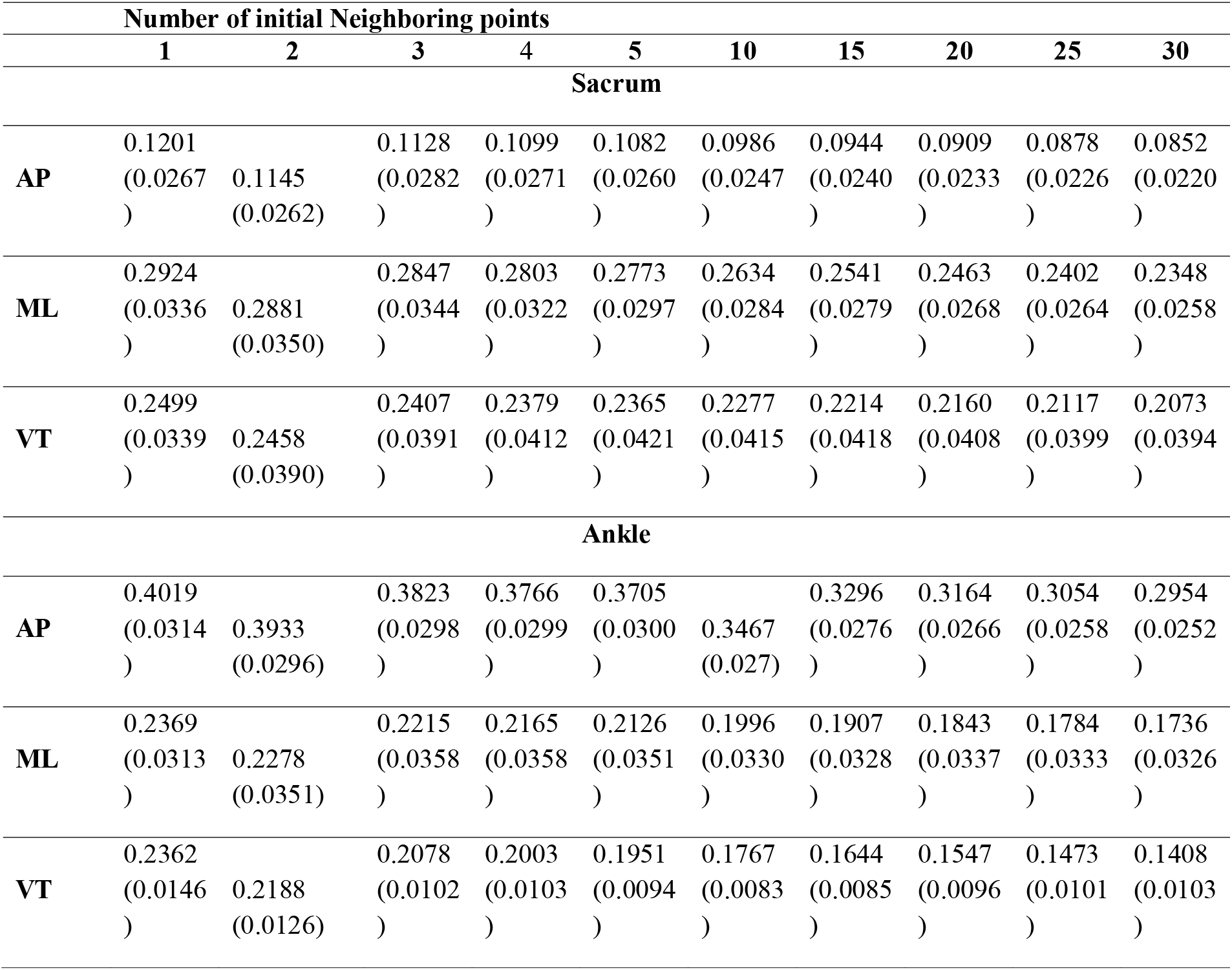
Average (standard deviation) of the largest Lyapunov exponent (LyE) calculated for the acceleration time series of sacrum and ankle acquired from accelerometer. The values are averaged over three trials. The 1-point neighboring condition is equivalent to the original Rosenstein’s method.

## Discussion

The objectives of this study were to investigate the effect of increasing the number of initial neighboring points on the LyE value calculated with Rosenstein’s method in the presence of noise as well as comparing the proposed method with the filtering method. The findings revealed that for both mathematical and experimental time series, the proposed method led to a consistent LyE value for neighboring points more than 15 data points. In addition, filtering with the residual frequency resulted in overestimation of LyE values compared to the original value as well as values calculated with the proposed method.

For almost all time series, the %Difference fell within 1% threshold from the 15-point neighboring condition onwards implying that selecting at least 15 initial neighboring points results in a reliable computation of LyE. This is also clear from the graphs (Figures 2 and 3) where they plateaued at the 15-point condition onwards. It should however be noted that this finding is specific to the current study where each cycle was time-normalized to 100 data points and the average mean period of the constructed state vectors was around 80 data points. Therefore, the emphasis of this study is to demonstrate the feasibility of the proposed method and not necessarily, the number of initial points found. Moreover, the 1% threshold found here was not set a priori and rather was a post hoc finding of the study. Therefore, in line with the previous point, the plateau threshold for biomedical signals with distinct characteristics could be different to what found in this study.

In addition, it was found that the %Difference was positive for mathematical time series whereas it was negative for experimental time series (Figures 2–4). This means the LyE value increased by increasing the number of neighboring points for simulated time series while it decreased for experimental time series. The reason lie in the difference between the added noise to simulated time series and the noise contained in the experimental time series. That is, by adding noise to a simulated time series, each data point on the trajectory is transferred (mapped) to a new point (noise data point) off the time series trajectory. Consequently, the average of initial distance of the neighboring trajectory is increased at each time step by increasing the number of neighboring points leading to increase of LyE value compared to the previous neighboring condition. For experimental time series on the other hand, the noise data point is separate to real time series data point. Thus, by increasing the number of neighboring points, more real data points are chosen which leads to a lower average initial distance at each time step and lower lyE values for higher neighboring conditions.

**Figure 4:**
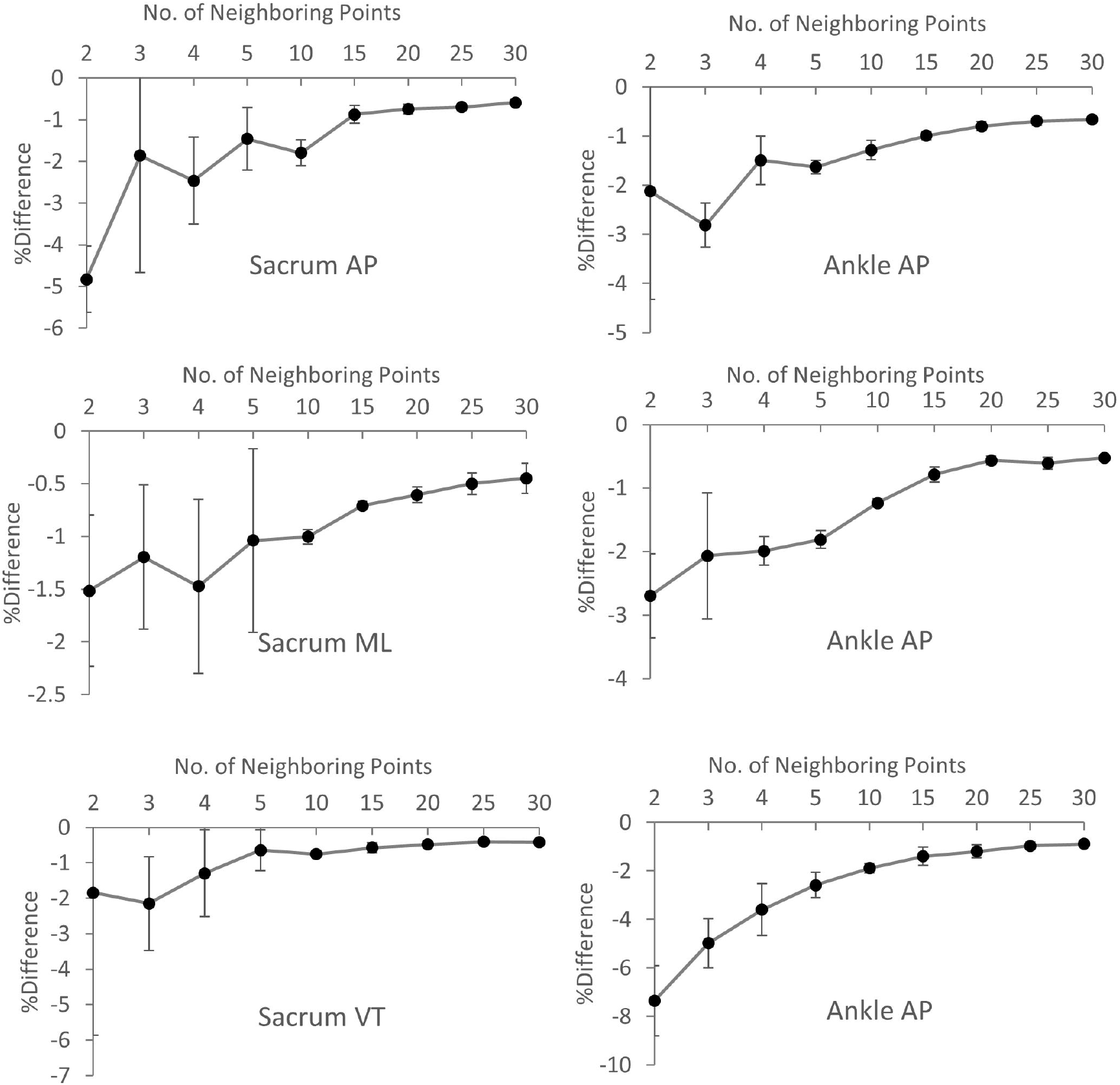
The %Difference calculated for sacrum (left) and ankle (right) acceleration time series in anterior-posterior (AP, top), medio-lateral (ML, middle), and vertical (VT, bottom) directions. The accelerometer was used to record segments’ acceleration. The 1-point neighboring condition is equivalent to the original Rosenstein’s method

The findings also indicated that a lowpass filtering with residual cutoff frequency resulted in overestimating of the LyE (Table 5). It should emphasized that the original Rosenstein method (i.e. only one neighboring point) was adopted to calculate the LyE for the filtered signals. This finding implies that using the revised Rosenstein’s method led to a more conservative value for LyE which at least has two advantages: (a) the LyE is in the range of original LyE revealing that, in contrast to the filtering method, the information contained in the time series is not missing, and (b) LyE is more consistent and more reliable than both original Rosenstein’s and filtering methods (see the plateau in the graphs). This could particularly be imperative when for instance, comparing time series variability between healthy individuals and patients. To explain, the inconsistent LyE values calculated with the original Rosenstein’s and/or filtering methods might not reveal the inherent nature of variability in the mentioned groups. Furthermore, the results of filtering in the current study was in line with the study of Mehdizadeh & Sanjari[13] where the filtered values of the LyE were higher than non-filtered values calculated using original Rosenstein’s method.

**Table 5:**
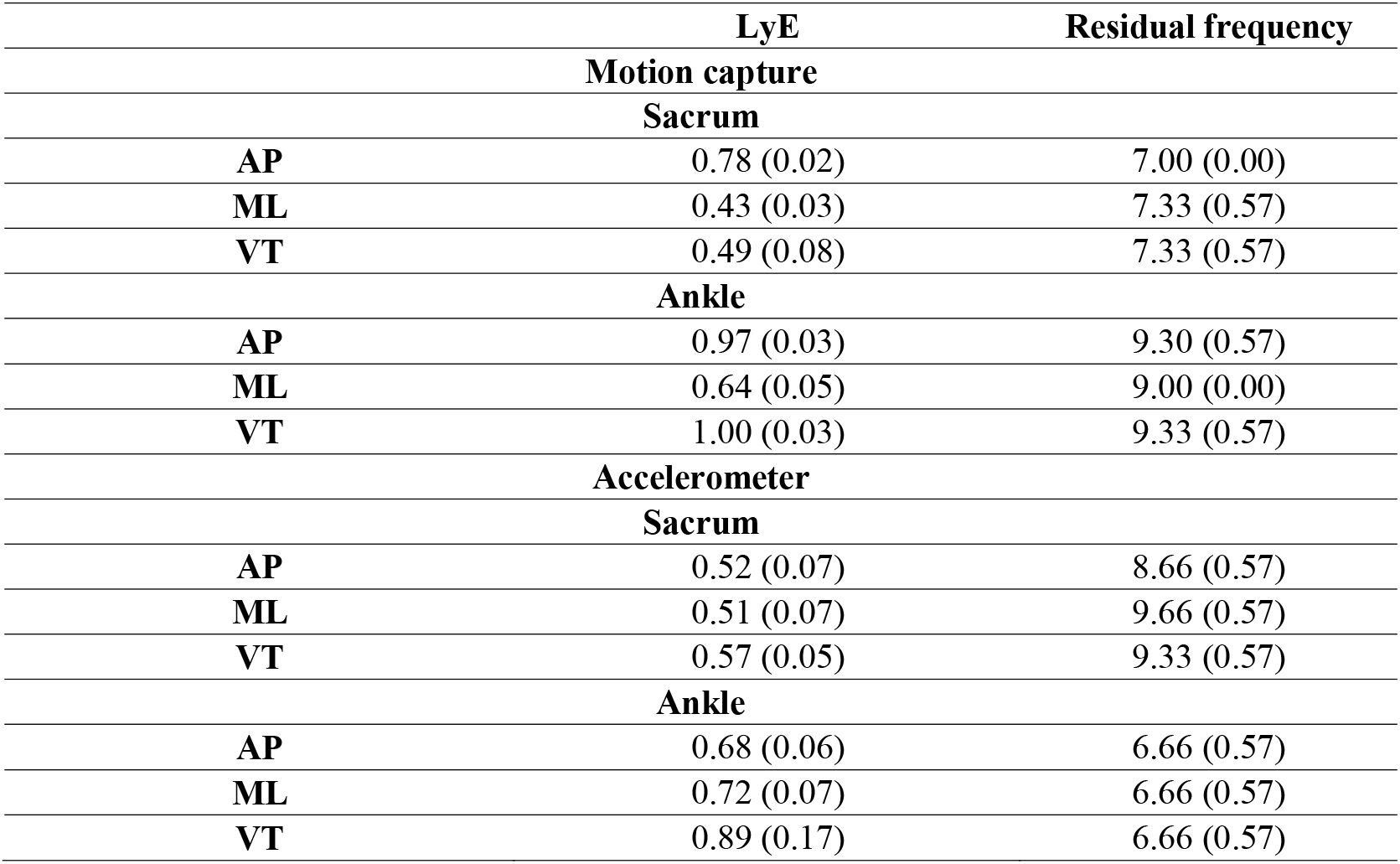
Average (standard deviation) of the largest Lyapunov exponent (LyE) calculated for the filtered time series of sacrum and ankle acquired from motion capture and accelerometer. The residual frequency used for the filtering is also depicted in the table. The values are averaged over three trials.

One point to mention here is that according to the original Rosenstein’s method, the distance of the neighboring point to the reference point should be greater than the mean period of the state vectors of the state space. In other words, the neighboring and reference points should not be on the same trajectory (i.e. cycle) in the state space. This also holds for the revised method where none of the neighboring points should be located in the distance less the mean period from the reference point. However, the neighboring points themselves should not necessarily be on different trajectories and can be within the same cycle. This will not violate the first assumption because the distance of each neighboring point to the reference point is greater than the mean period of the state space (Figure 1).

A note of caution is due here since this study should be viewed as a proof of concept study, the experiments were done on one healthy participant. Future studies could compare the revised Rosenstein method proposed in this study and the original Rosenstein’s method in larger groups of participants. The rationale being that since the new method yields to a more consistent and more reliable LyE value, it could better reveal the difference between healthy and non-healthy groups. In addition, the highest %Difference found for the experimental time series was in the range of 2-7% (the 2-point neighboring condition in Figures 3 & 4) which is not high compared to the plateau region (around 1%). Therefore, it might be argued that the proposed method is not efficient in terms of removing the effect of noise. However, as pointed out earlier, the emphasis of this study is not on the values calculated, but rather on the feasibility of the proposed method instead of the original Rosenstein’s method and the rationale behind it. In other words, the values are specific to this study and not generalizable. Using the proposed method in other studies might thus lead to different values compared to the current study.

In conclusion, this study proposed a revision to the Rosenstein’s method of calculating LyE to make it more robust to noise. This new method takes the advantage of choosing multiple neighboring points (rather than only one point as in the Rosenstein’s original method) at each step of computing divergence. The effect of noise could thus be canceled out by averaging over several neighboring points. The results of applying this method to both simulated and experimental time series demonstrated that the LyE graph reached a plateau at the 15-point neighboring condition inferring that the LyE values calculated using at least 15 neighboring points were consistent and reliable. Notwithstanding the relatively limited sample, the proposed method could be used to calculate LyE more reliably in experimental studies where noise is omnipresent.

## Acknowledgement

The tests for this study were done in the Biomechanics Lab at the National Sports Institute of Malaysia.

## Competing interests

No conflict of interest

## Funding

None

## Ethical approval

By Local organizational committee.

